# Allogeneic testes transplanted into partially castrated adult medaka (*Oryzias latipes*) can produce donor-derived offspring by natural mating over a prolonged period

**DOI:** 10.1101/2022.05.07.490875

**Authors:** Daichi Kayo, Shinji Kanda, Kataaki Okubo

**Author notes:** corresponding author Daichi Kayo.

## Abstract

Generally, successful testis transplantation has been considered to require immune suppression in the recipient to avoid rejection of the transplanted tissue. In the present study, we demonstrate in medaka that allogeneic adult testicular tissue will engraft in adult recipients immediately after partial castration without the use of immunosuppressive drugs. The allografted testes are retained in the recipient’s body for at least three months and are able to produce viable sperm that yield offspring after natural mating. Some recipients showed a high frequency (over 60%) of offspring derived from spermatozoa produced by the transplanted testicular tissue. Histological analyses showed that allografted testicular tissues included both germ cells and somatic cells that had established within an immunocompetent recipient testis. The relative simplicity of this testis transplantation approach will benefit investigations of the basic processes of reproductive immunology and will improve the technique of gonadal tissue transplantation.

## Background

Gonadal or germline transplantations have been used to investigations of reproductive biology/immunology and have also been successfully applied for selective breeding in livestock and aquaculture, species conservation, and fertility treatment. A variety of allogeneic or xenogeneic transplantation protocols for gonadal tissues or germ cells have been developed and used to create potentially superior broodstocks, as insurance against the accidental death of vital broodstocks and for maintenance of threatened breeds and species (1-6). One of the major drawbacks of allogeneic transplantation of tissues, however, is the possibility of immunorejection of the donor cells and tissues. The use of spermatogonial stem cells (SSCs) for transplantation is considered particularly valuable as these cells are present in large numbers in the testes of adult males and are relatively easy to obtain (7, 8). SSC transplantation studies in mice and rats have found that the donors and recipients need to be closely related to avoid a immunorejection; alternatively, immunodeficient animals can be used as recipients, or the recipients can be treated with immunosuppressant drugs (9, 10). In teleosts, the immunorejection problem can be avoided by transplanting germ cells into newly hatched recipient larvae whose immune systems are immature (11-14). However, this approach is technically demanding and requires the use of microinjection equipment.

Testis allografting is a possible alternative approach for germline transplantation, which can be easily performed, but the potential for immunorejection of donor cells and tissues remains with this method. A few sites in the body display “immune privilege” in which an antigenic response is not elicited by the presence of transplanted cells. The testes are known to have immune privilege and are more likely to accept transplanted tissues (immune privilege site) and also to be the source of donor cells (immune privilege tissue) (15). A similar phenomenon may exist in fish because it has been reported in fish that transplants of body tissue (scales) are rejected within a few days to about two weeks, while subcutaneous transplants of testicular tissue are accepted for six to nine weeks (16-18).

The present study was initiated to develop a reliable method for allogeneic testis transplantation in fish. We chose the model fish species medaka (*Oryzias latipes*) for our analyses, as they spawn daily, are amenable to gene editing, and a surgical method for gonadectomy has been established (19, 20). We demonstrated the immunocompetency of the recipient medaka used in the present study by scale transplantation experiments. However, as described above, the testis is immune privileged and histological analyses of recipient testes after transplantation showed that they contained donor germ cells and somatic cells. These results indicate the feasibility of developing a reliable method for creating male surrogate parents to efficiently obtain donor-derived offspring.

## Materials and methods

### Animals

All medaka used in the study were maintained under a 14 hour light/ 10 hour dark photoperiod (light from 09:00 to 23:00), with a water temperature of 28°C. The fish were fed 3–4 times per day with live brine shrimp (*Artemia nauplii*) and a commercial pellet food (Otohime; Marubeni Nisshin Feed, Tokyo, Japan). We used d-rR/TOKYO (d-rR) strain medaka, along with transgenic strains, and captive-bred wild-type medaka. Transgenic medaka that express GFP under the neuropeptide B promoter (*npba*-GFP) were used (21). Transgenic medaka consistently expressing GFP (strain ID: TG862, d-rR-Tg(beta-actin-loxP-GFP); *actb*-GFP) were obtained from the National Institute for Basic Biology via The National BioResouce Project-Medaka (NBRP-medaka). Please note that the d-rR strain is not an inbred strain. Thus, the *actb*-GFP medaka used as donor and recipient d-rR medaka are not isogenic with each other. Because *actb*-GFP strain females showed low fecundity, we generated the F1 hybrid (*actb*-GFP hetero) between *actb*-GFP strain males and recipient strain (d-rR) female, and *actb*-GFP hetero males were used as donor fish in some analyses. The ancestor of the wild-derived medaka was caught in an irrigation channel of a rice field (GPS coordinates: 32°58’21.9”N 132°58’12.6”E (32.972750, 132.970167); Isawa, Shimanto City, Kochi Prefecture). This wild-derived strain has been bred and maintained for a number of generations in our laboratory.

### Testis transplantation into recipient males

Medaka aged 3–8 months for each strain were used as donors; they were anesthetized, decapitated, and the testes were dissected. Isolated testes were kept in phosphate-buffered saline (PBS) until transplantation. Twenty-two recipient medaka (d-rR strain, aged 2–5 months) were anesthetized using 0.02% MS-222 and their abdomens were incised using a razor blade. In male medaka, the testis is essentially a single organ following the fusion of bilateral testes during ontogeny (22). The rostral side of the recipient testis was pinched using forceps, and most of the testicular tissue was removed, leaving a part of the caudal side of testis using another set of forceps. The isolated donor testis was cut into 1– 2 mm pieces which were placed adjacent to the remaining part of the recipient testis. After implantation, the abdominal incision was sutured with nylon thread. Post-surgical recovery was carried out by placing the recipient medaka in the 0.8% saline for two or three days; the fish were transferred to a freshwater environment after recovery. The abdomens of the recipient medaka and of their offspring were photographed using a stereomicroscope (M165FC or M205FA, Leica Microsystems, Wetzlar, Germany) equipped with a DFC7000T digital camera (Leica Microsystems). GFP fluorescence was detected using an excitation spectrum of 450–490 nm and emission spectrum of 500–550 nm.

### Scale transplantation experiments

The immune responses of the fish strains used were confirmed by scale transplantation experiments; *actb*-GFP strain, *actb*-GFP hetero, and wild-derived strain (6–7 months old) were used as the donor strains, and d-rR strain medaka (6–7 months old) were used as the recipients. As a control, we transplanted scales between siblings of the d-rR strain (4–5 months old) that had been maintained for a number of generations in our laboratory and, essentially, has the same genetic background, to confirm that body tissue transplants were not rejected by the immune system of these fish.

Four recipient medaka were anesthetized using 0.02% MS-222. A few donor medaka were anesthetized and decapitated; 20–23 scales were removed from the donor body and transplanted into the caudal region around the lateral line of the four recipients (Day 0). The recipients were kept in a tank throughout the experimental period. The number of engrafted scales on the recipients was counted each day and the fish were photographed on Days 1, 7, and 10 under an M205FA stereo microscope equipped with a DFC7000T digital camera. Fluorescent staining was viewed after 450–490 nm and 540–580 nm excitation and 500–550 nm 593–667 nm emission for GFP and Alizarin red S (ARS), respectively.

### Vital staining of scales

In the control analysis using d-rR siblings, we stained the scales of donor fish with ARS (Wako, Osaka, Japan), a vital stain for fishbone (23), to distinguish them from the scales of the recipient. Medaka were anesthetized using 0.02% MS-222 and dried with tissue paper. A saturated solution of ARS (0.1% ARS in PBS) was dropped onto the fish body with a micropipette and left for 10–60 seconds. Medaka with red scales were released into the tank and used as donors on the following day. Scale transplantation was performed as described above. The stained scales transplanted into recipients could generally be distinguished from the unstained scales of the recipient by eye for up to 5 days; after 6 days, it was necessary to use fluorescence to identify donor scales.

### Immunohistochemistry (IHC)

The testis of *actb*-GFP hetero (age 4–5 months, n = 2), recipient strain (age 4–5 months, n = 2), and a recipient that had been transplanted with a testis from an *actb*-GFP strain (age 6–7 months) or *actb*-GFP hetero fish were excised (n = 3, 16 days or 2 months after surgery) and fixed in Bouin’s fixative solution or 4% paraformaldehyde (PFA)/PBS. The fixed testis was dehydrated through an ethanol series, cleared with xylene, and embedded in paraffin. 10-μm sections were cut and treated with 0.3% H_2_O_2_ for 30 min, 2% normal goat serum (NGS) for 30 min, and incubated with anti-GFP rabbit polyclonal antibody (#598, Medical and Biological Laboratories, Tokyo, Japan) diluted at 1:500– 1:2000 in PBS containing 2% NGS overnight at 4°C. After two washes in PBS, the sections were incubated with biotinylated goat anti-rabbit IgG (diluted according to the manufacturer’s protocol) for 1 hour and stained using the VECTASTAIN Elite ABC reagent (VECTASTAIN(R) Elite ABC-HRP Kit, Peroxidase, PK-6101; Vector Laboratories, Burlingame, CA) for 1 hour. The horseradish peroxidase-conjugated Avidin-Biotin Complex was visualized using TSA Plus Fluorescein System (PerkinElmer, Waltham, MA, USA) or 3,3-diaminobenzidine (DAB) and 0.003% H_2_O_2_. Cell nuclei were counterstained with 4′,6-diamidino-2-phenylindole (DAPI) or hematoxylin. Fluorescent images were acquired by using a confocal laser scanning microscope (Leica TCS SP8; Leica Microsystems, Wetzlar, Germany). The following excitation and emission wavelengths, respectively, were used for detection: DAPI, 405 nm and 410–480 nm; fluorescein and Alexa Fluor 488, 488 nm and 495–545 nm.

### Dual labelling for GFP and mRNA of Sertoli/Leydig cell marker genes

To examine the co-existence of GFP and Sertoli/Leydig cell marker genes, we performed dual labelling for IHC and *in situ* hybridization (ISH) analysis. The testis of a recipient that had been transplanted with a testis from an *actb*-GFP strain or *actb*-GFP hetero fish was excised, fixed in 4% PFA/PBS for 4–6 hours, and embedded in paraffin (n = 2, 16 days after surgery). 10-μm sections were cut and hybridized with digoxigenin (DIG)-labeled RNA probe. The DNA fragments of *gsdf* (AB525390) as a Sertoli cell marker and *hsd3b* (AB525390) as a Leydig cell marker were used to generate DIG-labeled probes. The DIG-labeled *gsdf* probe was visualized by using an anti-DIG mouse primary antibody (Abcam, Cambridge, UK) and Alexa Fluor 555-conjugated goat anti-mouse IgG secondary antibody (Thermo Fisher Scientific, Waltham, MA, USA) while GFP was detected using an anti-GFP rabbit polyclonal antibody (Medical and Biological Laboratories), VECTASTAIN Elite ABC reagent (Vector laboratories), and TSA Plus Fluorescein System (PerkinElmer). The DIG-labeled *hsd3b* probe was visualized by using a horseradish peroxidase-conjugated anti-DIG antibody (Roche Diagnostics, Basel, Switzerland) and TSA Plus Cy3 System (PerkinElmer) while GFP was detected using an anti-GFP rabbit polyclonal primary antibody and an Alexa Fluor 488-conjugated goat anti-rabbit IgG secondary antibody (Thermo Fisher Scientific). Cell nuclei were counterstained with DAPI. Fluorescent images were acquired by using a confocal laser scanning microscope (Leica TCS SP8). The following excitation and emission wavelengths, respectively, were used for detection: DAPI, 405 nm and 410–480 nm; fluorescein and Alexa Fluor 488, 488 nm and 495–545 nm; and Cy3 and Alexa Fluor 555, 552 nm and 562–700 nm.

## Results

### Adult donor testis transplanted into an adult recipient male is functionally engrafted without immunosuppression

We performed testis transplantation using *actb*-GFP donors and d-rR recipients. Four of the ten d-rR males whose testis was partially replaced with an *actb*-GFP testis showed strong green fluorescence in their abdomens at 2 months after surgery (Figure 1 a–c). Thus, successful allografts were present in four of the fish. To determine whether the engrafted testis was functional, we mated the GFP-positive recipients with d-rR females and assessed the frequency of GFP-positive eggs 2-7 weeks after surgery (Figure 1 d, e; Table 1). The frequency of GFP-positive eggs was approximately 9, 18, and 66% for three fish; the fourth fish produced no GFP positive eggs (Table 1). We also performed testis transplantation using donor *npba*-GFP medaka that were generated in our laboratory and had the same genetic background as the recipient fish (Table 1, #5 and #6). Two of the four recipients had high frequencies (95% and 100%, respectively) of GFP-positive eggs (Table 1). These results demonstrated that an adult testis allografted into an adult recipient male is functional.

**Table 1.** Results of the mating analysis: surrogate father of d-rR strain allografted with *actb*-GFP strain or *npba*-GFP strain testis.

| After surgery (weeks) | 2 to 7 | 2 to 7 | 2 to 7 | 2 to 7 | 2 | 2 to 7 |
| --- | --- | --- | --- | --- | --- | --- |
| Individual | #1 | #2 | #3 | #4 | #5 | #6 |
| GFP + | 3 | 70 | 9 | 0 | 20 | 54 |
| GFP - | 30 | 36 | 40 | 78 | 1 | 0 |
| % | 9.09 | 66.04 | 18.37 | 0.00 | 95.24 | 100.00 |

GFP-positive eggs, which indicates fertilization by sperm from the allogenic or isogenic donor testis, were produced by four of the ten (individuals #1~#4) or two of the four (individuals #5 and #6) recipient males, respectively.

**Figure 1.**
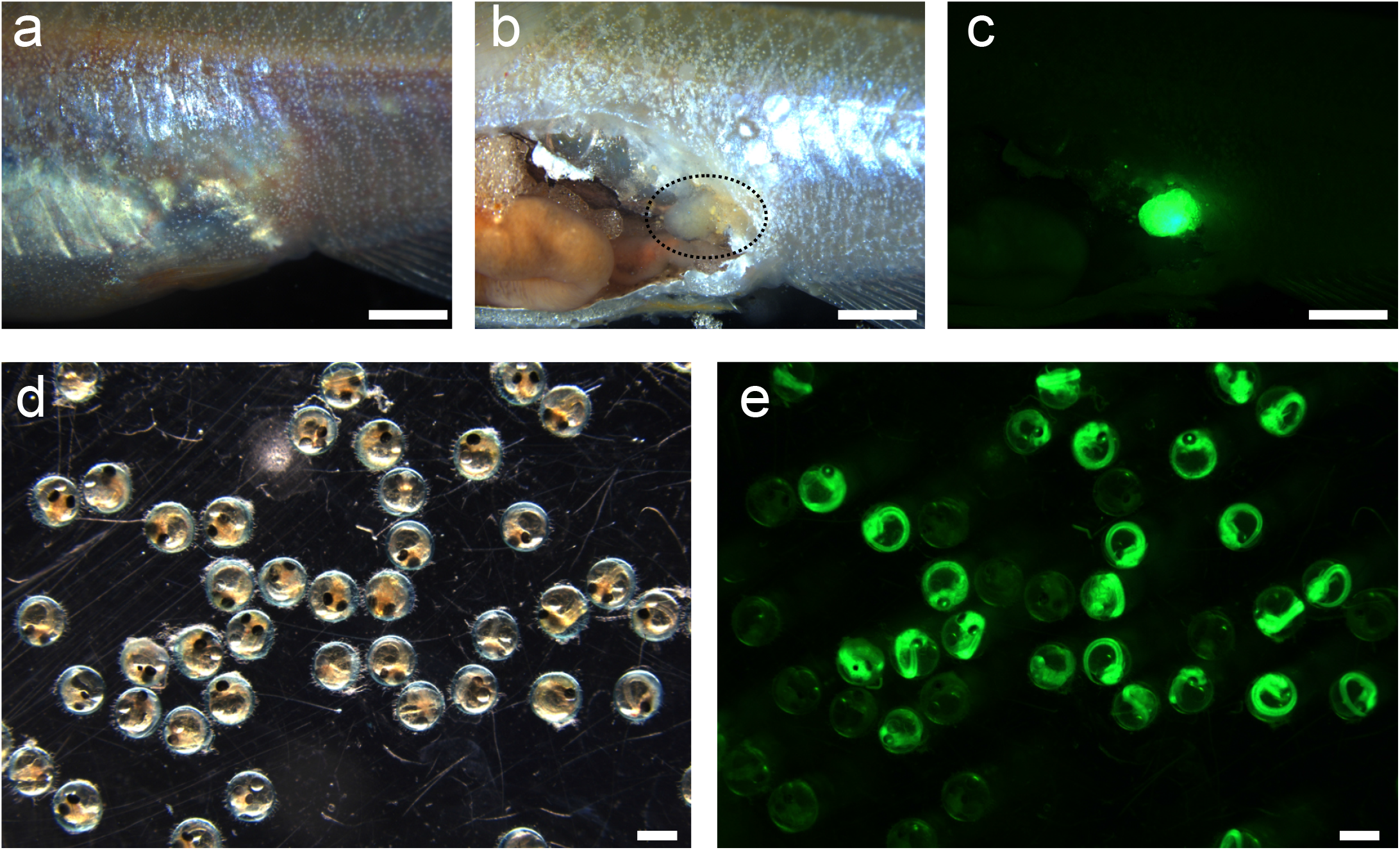
The appearance and functionality of testicular tissue allografted into the abdomen of a recipient. (a–c) Representative images of a recipient male that received testicular tissue derived from an *actb*-GFP strain male. Bright field image of the intact abdomen (a), bright field image of the incised abdomen (b), and fluorescence image of the incised abdomen (c); transplanted location is encircled by a dotted line in panel (b); scale bar, 1 mm. (d, e) Representative images of the eggs fertilized by the recipient male that had been transplanted with testicular tissue of an *actb*-GFP strain male. Bright field image (d) and fluorescence image (e); scale bar, 1 mm

### Functional allografts produced by transplanting testis from wild-derived medaka into d-rR recipients

To determine whether testis transplantation can be applied to genetically distant strains, we transplanted testes from wild-derived medaka into d-rR strain medaka males. The wild-derived medaka strain belongs to a different subclade than the d-rR strain due to geographical isolation (24) and has black pigmented scales. We also allografted testes from wild-derived strain donors to d-rR male recipients (Figure 2 a). Testicular tissues from wild-derived males were transplanted into eight d-rR males; the recipients were subsequently mated with d-rR females (Figure 2 a, b). Interestingly, black pigmented eggs, which indicate fertilization by sperm from the wild-derived donor testis, were produced by two of the eight recipients (Figure 2 c). All the fertilized eggs of one of these recipients (#7) were pigmented; the other produced 9% pigmented eggs (Table 2). These results showed that the testis transplantation was feasible even if the donor’s genetic background was distant to the recipient (d-rR) strain.

**Table 2.**
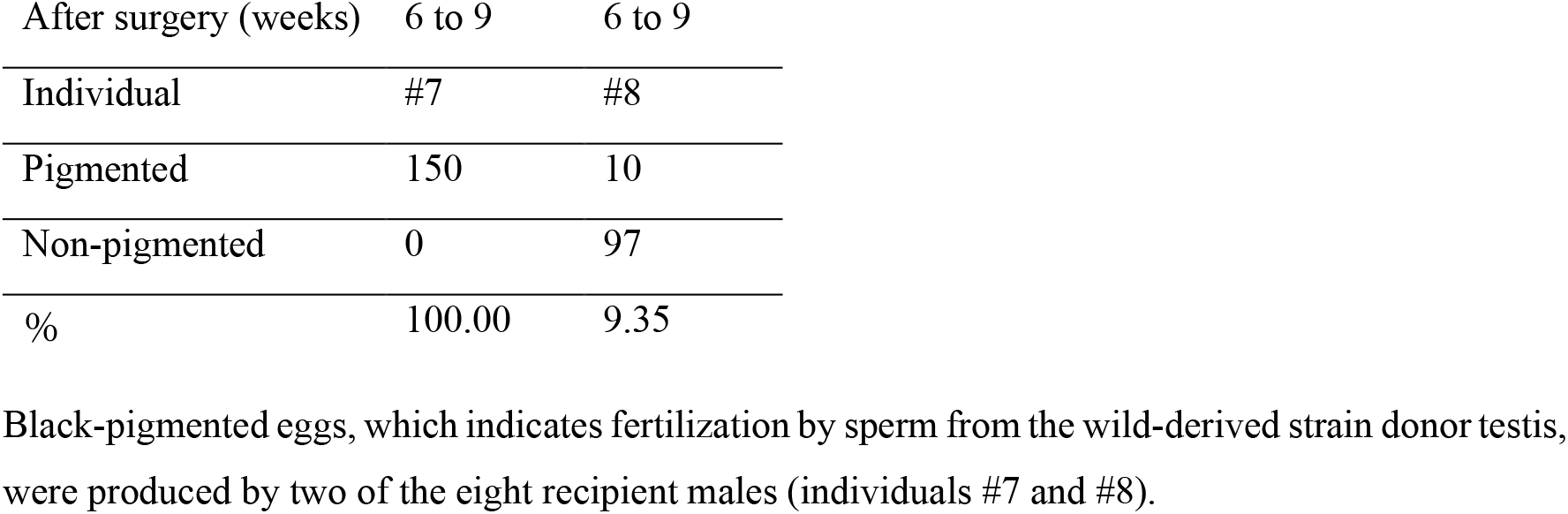
Results of the mating analysis: surrogate father of d-rR strain allografted with wild-derived strain testis.

| After surgery (weeks) | 6 to 9 | 6 to 9 |
| --- | --- | --- |
| Individual | #7 | #8 |
| Pigmented | 150 | 10 |
| Non-pigmented | 0 | 97 |
| % | 100.00 | 9.35 |

Black-pigmented eggs, which indicates fertilization by sperm from the wild-derived strain donor testis, were produced by two of the eight recipient males (individuals #7 and #8).

**Figure 2.**
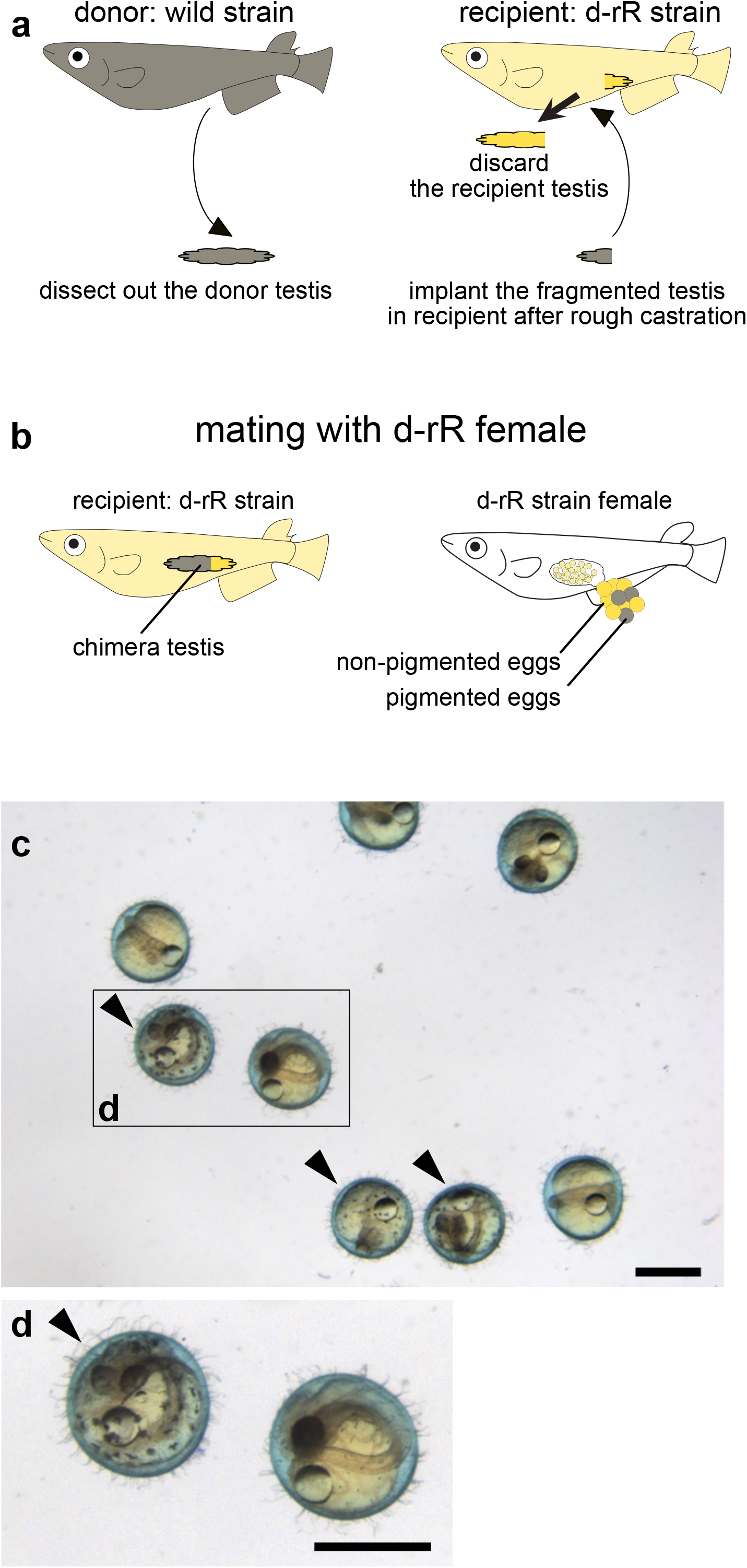
Functional allografts of testicular tissue from donor medaka with a different genetic background to the recipients. (a) An outline of the surgical procedure used here. (b) An outline of the mating scheme used here. In medaka, females lay eggs after spawning and keep the eggs attached to their belly for a while. Pigmented eggs are produced following fertilization by spermatozoa of wild-derived strain germ cells. Non-pigmented eggs result from fertilization with d-rR strain sperm. (c) Representative image of eggs fertilized by a recipient that had been transplanted with testicular tissue from a wild-derived medaka strain; arrowhead, pigmented egg resulting from fertilization with a wild-derived spermatozoon; scale bar, 1 mm. The boxed area is magnified in panel (d).

### Transplanted scales are rejected by the immune system of the recipient

We performed a scale transplantation experiment to confirm that d-rR recipients would reject somatic tissues from other strains (Figure 3 a–f, and Table 3). Loss of transplanted scales may be caused by immunorejection or mechanical injury; these two causes can be distinguished by the fact that mechanical injury during the transplantation process results in the loss of the scales on the day after transplantation (18). Our analysis of the recipient fish on successive days after scale transplantation indicated that 10–15 scales derived from wild-derived and *actb*-GFP strain fish had been engrafted into recipients. Almost all the transplanted scales were rejected by days 7 to 9, and all scales were lost within 12 days. To confirm that the scale transplantation was successful, we performed vital staining of the scales with ARS in d-rR donors and transplanted these stained scales into d-rR recipients (Figure 3 g, h). After the loss of some scales on Day 1 due to mechanical injury, most of the allografted d-rR scales had been accepted at 12 days by the d-rR recipient (Table 3). *actb*-GFP strain were generated from the d-rR strain, and therefore their genetic backgrounds should be the same. However, it should be noted that the d-rR strain is not an inbred strain. Based on the fact that transplanted *actb*-GFP scales were rejected by the recipient immune system, we conclude that the genetic backgrounds are sufficiently distant to cause immunorejection. Our results demonstrate that recipient d-rR strain medaka reject allografted tissues from donor medaka (*actb*-GFP strain and wild-derived strain).

**Table 3.** Results of scale transplantation into a d-rR recipient.

|  |  | Day0 | Day1 | Day2 | Day3 | Day4 | Day5 | Day6 | Day7 | Day8 | Day9 | Day10 | Day12 | Day17 |
| --- | --- | --- | --- | --- | --- | --- | --- | --- | --- | --- | --- | --- | --- | --- |
| **Pigmented scale** | observed | 23 | 10 | 10 | 9 | 9 | 8 | 8 | 6 | 6 | 0 | - | - | - |
|  | lost |  | 13 | 0 | 1 | 0 | 1 | 0 | 2 | 0 | 6 | - | - | - |
| **GFP scale** | observed | 23 | 15 | 15 | 15 | 10 | 7 | 4 | 3 | 2 | 1 | 1 | 0 | - |
|  | lost |  | 8 | 0 | 0 | 5 | 2 | 3 | 1 | 1 | 1 | 0 | 1 | - |
| **Alizarin red S positive scale** | observed | 20 | 13 | 13 | 13 | 13 | 13 | 13 | 12 | 12 | 12 | 12 | 12 | 12 |
|  | lost |  | 7 | 0 | 0 | 0 | 0 | 0 | 1 | 0 | 0 | 0 | 0 | 0 |

**Figure 3.**
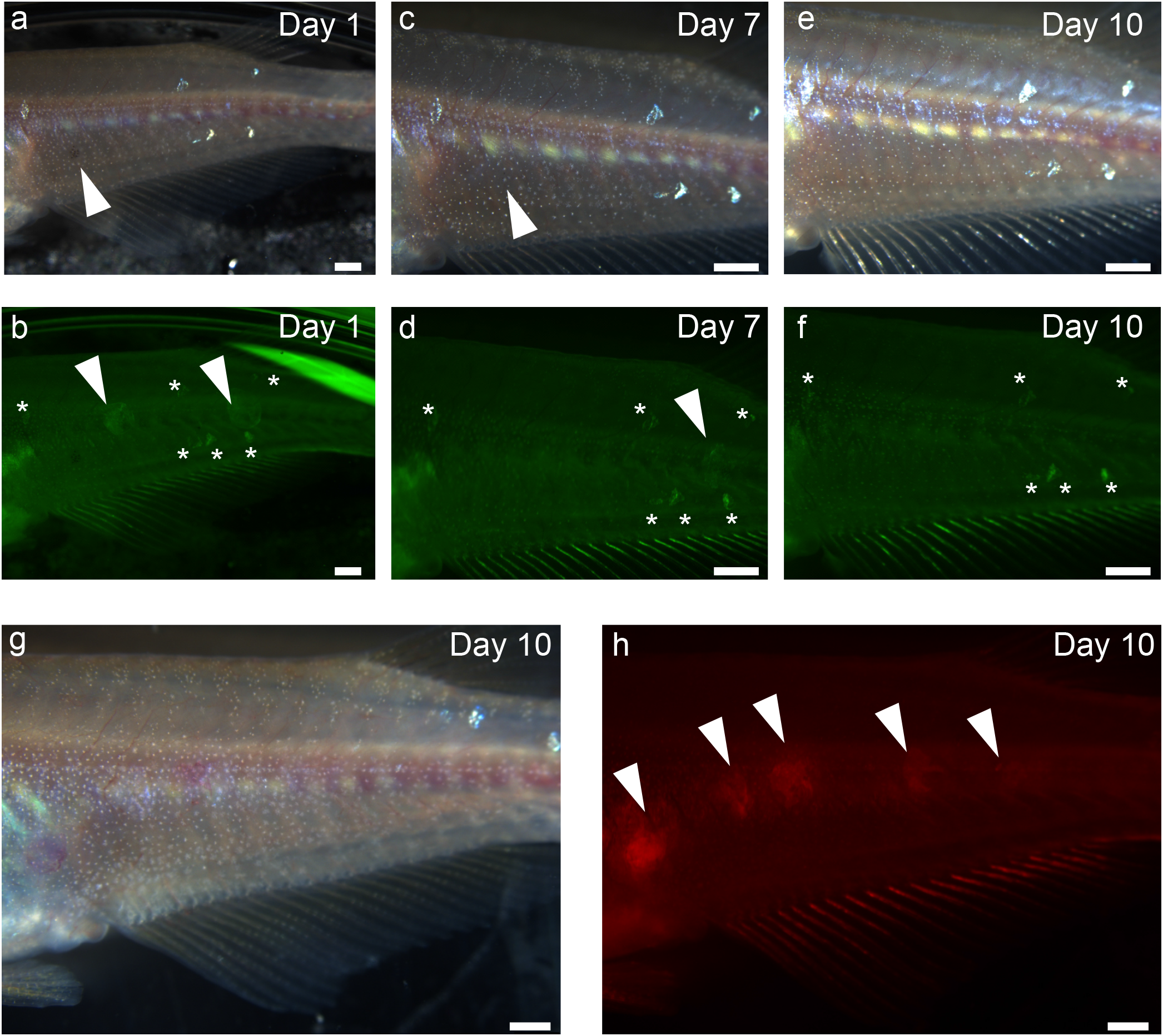
Allografted scales were immunologically rejected by the recipient. Representative images of transplanted allogenic (a–f) or isogenic (g, h) scales into a recipient. (a, c, e). Representative bright field images of scales from a donor (*actb*-GFP strain and wild-derived strain) transplanted into a recipient (d-rR strain). Day 1 (a), Day 7 (c), Day 10 (e); arrowhead indicates transplanted scale. (b, d, f) Representative fluorescence images of scales transplanted into a recipient. Day 1 (b), Day 7 (d), Day 10 (f); arrowhead indicates transplanted scale. Asterisk, autofluorescence originating from a scale on the recipient. (g, h) Representative images of a donor (d-rR strain) whose scales were vital stained with ARS and transplanted into a d-rR strain recipient: bright field image (g), and fluorescence image (h) on Day 10. Arrowhead, transplanted scale. Scale bar, 1 mm

### Allografted testes are functionally retained in recipients for more than 3 months

To determine the functional longevity of donor-derived testis in recipient medaka, we mated recipients for up to 13 weeks after surgery (Table 4). One recipient (#2) was sacrificed for abdominal analysis, and a second (#5) died accidentally; the other recipients were included in this analysis. As described in Table 4, four individuals (#3, #6, #7, and #8) showed almost equal frequencies of donor-derived eggs; two males did not produce any donor-derived offspring (Table 1); they are described as #1 or #4 in Table 4. This analysis demonstrated that allografted testis remained functional over an extended period of at least 13 weeks, except for one individual.

**Table 4.** Results of the mating analysis at 13 weeks or more after surgery: surrogate father of d-rR strain allografted with *actb*-GFP strain or wild-derived strain testis.

|  |  |  |  |  |  |  |
| --- | --- | --- | --- | --- | --- | --- |
| After surgery (weeks) | 13 | 13 | 13 | 15 | 13 | 13 |
| Individual | #1 or #4* | #1 or #4* | #3 | #6 | #7 | #8 |
| GFP + or pigmented eggs | 0 | 0 | 15 | 97 | 86 | 20 |
| GFP - or non-pigmented eggs | 94 | 17 | 69 | 0 | 0 | 106 |
| % | 0.00 | 0.00 | 17.86 | 100.00 | 100.00 | 15.87 |

Males of d-rR strain were used as recipients. The genetic backgrounds of donor testis were as follows; #1~#4, *actb*-GFP; #6, *npba*-GFP; #7 and #8, wild-derived strain. \*, #1 or #4 could not be distinguished.

### Male germ cells and somatic cells derived from the donor testis engraft into recipient testis

We performed an IHC analysis to detect GFP-expressing cells derived from the donor testis. GFP-positive cells (donor-derived cells) were distinguished as DAB-positive cells in histological sections, while GFP-negative cells (recipient cells) were only stained with hematoxylin (Figure 4 a–d). We used the *actb*-GFP strain and *actb*-GFP hetero medaka as donor males for the histological analysis. To confirm the immune rejection of the *actb*-GFP hetero donor in the recipient, we performed a scale transplantation analysis and demonstrated the immunocompetence to the donor scales in the recipient medaka (Table 5). All scales were rejected within 16 days.

**Table 5.**
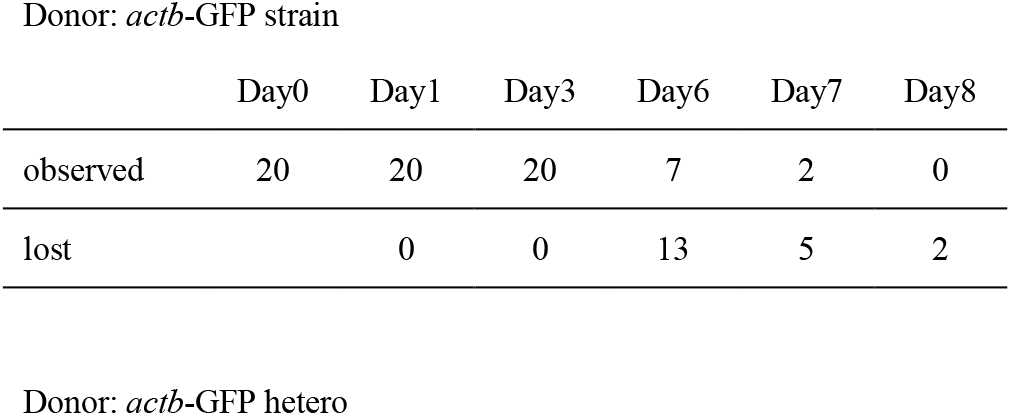
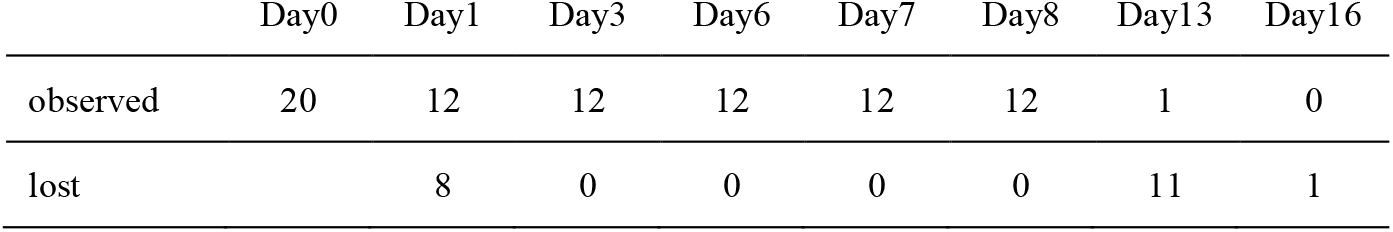
Results of *actb*-GFP medaka and *actb*-GFP hetero scale transplantation into a d-rR recipient.

**Figure 4.**
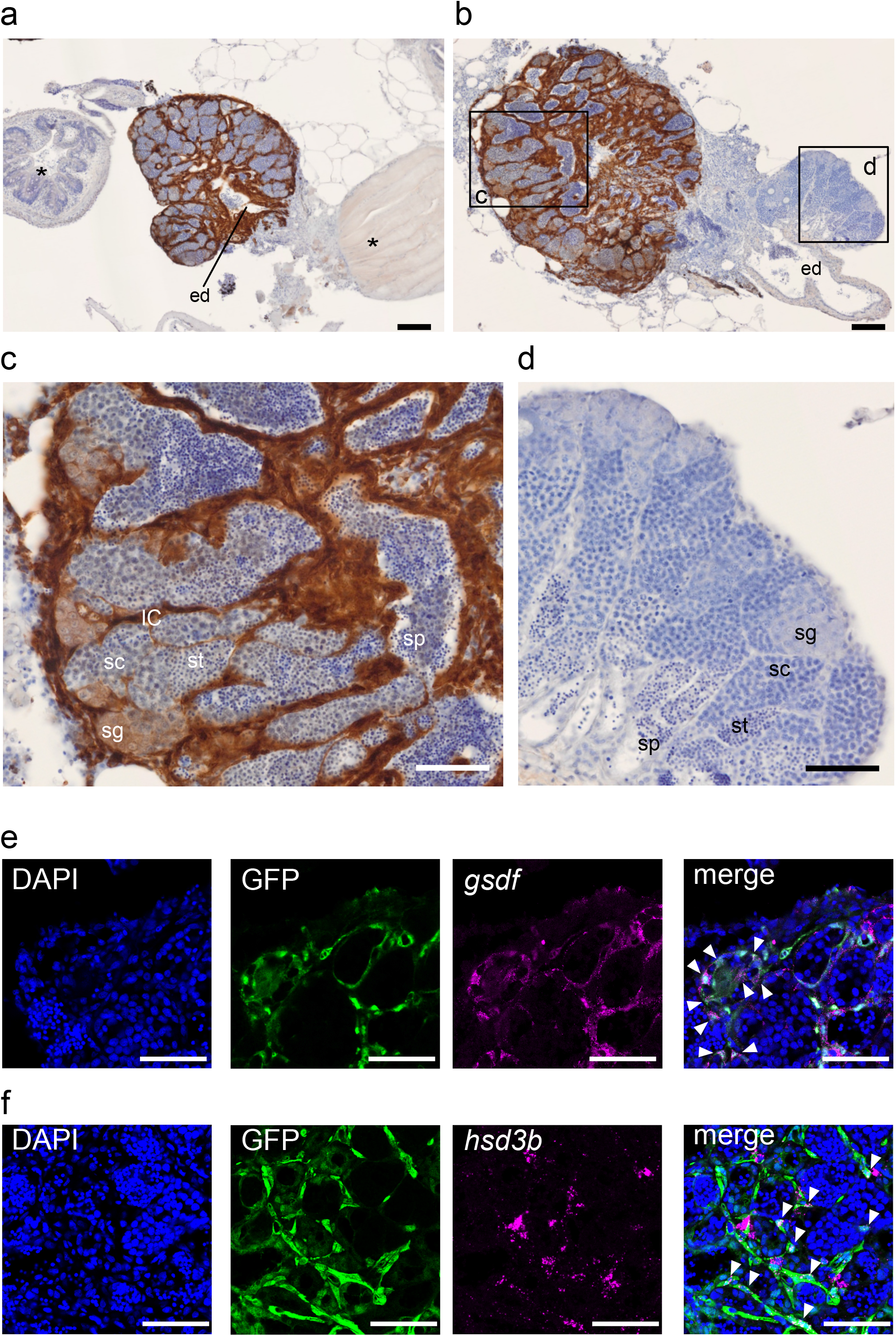
Allografted testicular tissue, including somatic cells, was fused with the recipient testis. (a–d) Representative images of IHC analysis using an anti-GFP antibody visualized by DAB. Sections were counterstained by hematoxylin. DAB-positive cells are donor-derived (*actb*-GFP strain) cells. (a) Representative image from IHC analysis of the recipient whose testis was mainly derived from allografted (donor-derived) testis. The *actb*-GFP strain was used as a donor male. ed, efferent duct. Asterisks denote non-gonadal tissue of the recipient. Scale bar, 100 μm. (b) Representative image from IHC analysis of the recipient whose testis was partly derived from allografted testis. F1 hybrid of *actb*-GFP strain and recipient strain (*actb*-GFP hetero) were used as a donor male. Scale bar, 100 μm. The boxed areas are magnified in panel (c): GFP-positive area (donor-derived tissue) and (d): GFP-negative area (recipient-derived tissue). IC, interstitial cells; sg, spermatogonia; sc, spermatocyte; st, spermatid; sp, spermatozoa. Scale bar, 50 μm. (e, f) Identification of the presence of Sertoli or Leydig cells in the allografted testis. The panels show the images of nuclear counterstaining (DAPI, blue), the cells of allografted testis (GFP, green), the expression of indicated marker genes ((e) *gsdf*, (f) *hsd3b*; magenta), and the merged image from the left in the same sections. Arrowheads denote representative cells that showed co-existence of the GFP and indicated marker genes. Scale bar, 50 μm.

For the classification of each developmental stage of spermatogenesis, we used the descriptions provided in previous studies (25, 26). GFP signals were detected in the allografted testis of the recipient male (Figure 4 a, b). The recipient testis contained spermatogonia with GFP signals, indicating that these spermatogonial cells proliferated and supplied the donor-derived germ cells. These observations also indicate the reason why some recipients produced donor-derived offspring over a long period. Interestingly, testicular somatic cells, such as interstitial cells (IC), had allografted into the recipient testis (Figure 4c). In our observations, donor-derived germ cells were surrounded by donor-derived somatic cells, not by recipient-derived somatic cells. These observations suggest that the donor-derived testicular tissue probably included Sertoli cells and Leydig cells that were not immunorejected but integrated into the recipient testis and supported functional spermatogenesis.

To analyze the presence of donor-derived Sertoli cells and Leydig cells after testis grafting, we performed dual labelling IHC/ISH analysis using the anti-GFP antibody and probes against *gsdf* as a Sertoli cells marker (27) and *hsd3b* as a Leydig cells marker (28). The expression of both marker genes was detected in the GFP-positive (donor-derived) cells in the allografted testis in the recipient male (Figure 4 e, f). These observations showed that both Sertoli cells and Leydig cells derived from allografted testis exist in the recipient male. We could scarcely detect fluorescent GFP signal in the allografted germ cells (Figure 4 e, f). Similar to this, the GFP signal of the germ cells was relatively weak compared to that of surrounding somatic cells in *actb*-GFP hetero male testis (Supplementary Figure 1). However, it was obvious that donor-derived germ cells exist in the allografted testis because we could get donor-derived offspring from recipient male (Figure 1 and 2). These results may suggest that the transcriptional activity of beta-actin is relatively low in germ cells.

## Discussion

In the present study, we demonstrated that transplanted allogeneic testicular tissue could engraft in the body of recipient adult medaka without the use of an immunosuppressive treatment. Additionally, we showed that allografted testicular tissue derived from medaka with a different genetic background was functional and produced sperm that resulted in fertilized eggs after natural mating. A histological analysis also showed that both germ cells and testicular somatic cells were engrafted into allogeneic adult recipients.

As some recipients fertilized eggs with donor-derived sperm by natural mating (Tables 1, 2, and 4), the sperm derived from the donor testicular tissue must have been released to the efferent duct, which was re-established after the transplantation surgery. From our histological observation, it seems that the genetic origin of the efferent duct is likely to be both donor- (Figure 4 a) and recipient-derived (Figure 4 b). It is interesting that the allografted testicular tissue, which included somatic cells, was accepted by the immunocompetent recipient whose genetic background was distant to that of the donor (Figure 2; Tables 2 and 4). In domesticated mammals, such as pigs and goats, it has been reported that allografted germ cells and Sertoli cells, successfully engraft in a recipient testis without the use of immunosuppressive treatment (3, 5). Our transplantation experiments here demonstrate that allogeneic transplantation of testicular tissue can succeed even in medaka with divergent genetic backgrounds. Examination of the geographic distribution of mitotypes of Japanese medaka (24) showed that the wild-derived medaka strain used as a donor in the present study belongs to subclade B-V, while the d-rR strain belongs to the subclade B-II; the divergence time among the B subclades is estimated as 0.5– 2.3 mya. These results suggest the feasibility of the present method for testis allografting, at least in medaka. However, because our results were obtained from a relatively low number of fish, the generality of our approach should be carefully interpreted.

Generally, allografted tissue is rejected by the immune system of the recipient. A previous study of allogeneic scale transplantation in medaka confirmed this expectation, as the allografted scales were rejected within 7 days (18). We confirmed that the recipient strain used here was immunocompetent by allografting scales from a wild-derived strain (black scales) and the *actb*-GFP strain into recipient d-rR strain fish; scales derived from the genetically distant donor were rejected within 12 days (Figure 3 and Table 3). Although the genetic backgrounds of the recipients (d-rR) and *actb*-GFP (generated from d-rR strain) might be expected to be similar, these strains are not inbred and have different genetic backgrounds. These results show that testicular tissue can engraft in allogeneic individuals, whereas somatic tissue, such as scales, are rejected by the immune system. This finding is consistent with the general consensus that testes have immune privilege (15). In a previous study on rainbow trout, testis allografted into subcutaneous tissue was retained for 6-9 weeks but rejected after 9 weeks (16, 17). In the present study, testicular allografts inserted into the abdomen of the recipient were retained for the full duration of our-13 week studying (Table 4). These results indicate that allografted testicular tissue is more readily accepted by the recipient than other somatic donor tissues.

In the present study, histological analyses were performed to analyze the cellular structure of the testicular allograft (Figure 4). Our results revealed that the allografted testis was fused with the recipient-derived testis. Here, we demonstrated that the donor-derived germ cells were surrounded by donor-derived somatic cells but not recipient-derived cells. In medaka, we occasionally observe the functional regeneration of testis after partial castration. According to a previous study, testicular tissue can regenerate functionally after partial castration in rainbow trout (29). Given this report and our observation, it is possible that the remaining part of the recipient testis was fused with donor-derived testicular tissue during the regeneration process.

GFP signals were observed not only in the germ line cells but also in the testicular somatic cells, such as the Sertoli cells and interstitial cells, which include blood vessels and Leydig cells (Figure 4 c) (30). Some of the testicular somatic cells (Sertoli and Leydig cells) are considered to play a role in immune tolerance in the testis. Sertoli cells create a local tolerogenic testicular environment in the testis by expressing immunoregulatory factors, such as serine protease inhibitor and clusterin, which down-regulate the signaling cascade under an antigen-antibody complex (31). Leydig cells, which produce sex steroid hormones in male testis, indirectly help the tolerogenic function of Sertoli cells by the actions of androgens (32, 33). Therefore, it is possible that donor-derived Sertoli and Leydig cells may assist allografted testis to evade the immunorejection by the recipient male. In contrast, an ovarian allografting study in rainbow trout demonstrated that allografted ovaries could not be accepted in other individuals (34). There might be also be a mechanism of immune tolerance that is regulated by these immune suppressive factors released from the testis in teleosts.

Methods for allogeneic or xenogeneic transplantation of SSCs, which are abundant in the testis, have been developed in many species. The methods for germ cell transplantation in teleosts can be classified into three approaches (35): primordial germ cell transplantation in fish embryos (36); germ cell transplantation in hatched fish larvae (12, 14, 37-39); and germ cell transplantation in adult fish (40-45). The latter method, germ cell transplantation in adult fish, has potential advantages over the other two approaches for aquaculture and species preservation. For example, it avoids the time lag between transplantation and sexual maturity of the recipient. Moreover, it does not require sophisticated techniques and equipment for microinjection into eggs or larvae. Adult tissue transplantation is relatively easy as it involves a simple transplantation procedure through the genital duct of the recipient after germ cell extraction from the donor testis (44, 45). To improve the success rate of germ cell transplantation to allogeneic individuals, it is considered crucial that the germ cells of the recipient are depleted but that the ability of the recipient to nurse donor-derived germ cells is maintained (1, 46, 47), e.g. through use of triploid individuals (48) or *dead end* gene knockdown fish (49, 50). Cytotoxic drugs such as busulfan may be used for germ cell depletion; use of these drugs adds a relatively short time to recipient preparation (2–4 weeks) (40, 42, 43). However, the study using cytotoxic drug reported that the frequency of offspring derived from donor sperm generally does not exceed 40% (44). In the present study, the method for germ cell transplantation is completely different from these studies because the testicular tissue is also allografted with male germ cells. Some of the recipients that had received donor testicular tissue immediately after partial castration showed a high rate (60–100%) of offspring derived from donor spermatozoa (Tables 1 and 4). This may be due to co-engraftment of germ cells and somatic cells in the transplanted testicular tissue, and the donor-derived testicular tissue may be able to nurse their own germ cells (Figure 4).

Cryopreservation methods for the whole testis have been developed in medaka (51). The combined use of testicular cryopreservation and the present approach for testicular tissue transplantation using adult recipients and natural mating may make it possible to shorten the time for recovery of larger numbers of offspring from cryopreserved testes compared to artificial insemination using cryopreserved sperm or injection of germ cells into larvae. In our IHC analysis, we observed GFP-positive spermatogonia (Figure 4 c). In medaka, it takes at least 5 days for spermatogonia to develop into spermatids and approximately one week for the spermatids to metamorphose into spermatozoa (52, 53). We mated each recipient used in the analysis here with three d-rR females for 2–3 weeks. Therefore, spermatogenesis in the donor-derived testis had sufficient time to complete at least one cycle of maturation before the mating analysis (Table 4). Our results suggest that the allografted germ cells proliferated in the recipient testis, allowing the recipient males to produce donor-derived offspring over a prolonged period (13–15 weeks). The rate of success for functional engraftment was approximately 30% in the present study; it will be necessary to improve this success rate to enable development of a simple, fast, and effective approach for testicular transplantation into adult recipient fish. It should also be noted that the present method requires the separation of donor-derived and recipient-derived offspring.

## Conclusions

We demonstrated the feasibility of allografting testicular tissue into immunocompetent recipients whose genetic background was distinctly different to those of the donors; functional engraftment was achieved after partial castration of the recipient without use of immunosuppressive treatments or chemical castration of the recipient. Further studies are required to improve our understanding of the immunological responses after testicular transplantation, and the results of these will be of value for aquaculture.

## Abbreviations

ARS: Alizarin red S,
DAB: 3,3-diaminobenzidine,
DAPI: 4′,6-diamidino-2-phenylindole,
ed: efferent duct,
DIG: digoxigenin,
IC: interstitial cells,
IHC: immunohistochemistry,
ISH: *in situ* hybridization,
MS-222: Ethyl 3-aminobenzoate methanesulfonic acid salt,
NGS: normal goat serum,
PBS: phosphate-buffered saline,
PFA: paraformaldehyde,
sc: spermatocyte,
sg: spermatogonia,
sp: spermatozoa,
SSC: spermatogonial stem cells,
st: spermatid

## Declaration of competing interest

### Ethics approval and consent to participate

All animal procedures were performed in accordance with the guidelines of the Institutional Animal Care and Use Committee of the University of Tokyo. The committee requests the submission of an animal-use protocol only for use of mammals, birds, and reptiles, in accordance with the Fundamental Guidelines for Proper Conduct of Animal Experiment and Related Activities in Academic Research Institutions under the jurisdiction of the Ministry of Education, Culture, Sports, Science and Technology of Japan (Ministry of Education, Culture, Sports, Science and Technology, Notice No. 71; June 1, 2006). Accordingly, we did not submit an animal-use protocol for this study, which used only teleost fish and thus did not require approval by the committee.

### Consent for publication

Not applicable

### Competing interests

The authors declare no conflict of interest.

### Availability of data and materials

All data generated or analyzed during this study are included in this published article.

## Funding

This work was supported by Grants-in-Aid from Japan Society for the Promotion of Science (JSPS) Grant 20K22587 to D.K. and NIBB Collaborative Research Program (22NIBB711) to D.K.

## Authors’ contributions

D.K. carried out all experimental work, designed the study, and drafted the manuscript; S.K. and K.O. helped in the interpretation of the data and preparation the manuscript. All authors gave final approval for publication.

## Acknowledgements

We thank Masato Kinoshita, National Institute for Basic Biology, and the National BioResource Project (NBRP) Medaka for providing the transgenic medaka (strain ID: TG862) in this study. We also thank Thomas Fleming for language editing of the revised manuscript.

## Figure legends

**Supplementary Figure 1. The protein level of GFP in the germ cells is relatively low compared to that of the surrounding somatic cells.**

(a, b) Representative images from the IHC analysis using an anti-GFP antibody visualized by DAB staining (a) or fluorescent detection (b). (a) Left panel shows the image of the testis that consistently express GFP with beta-actin (*actb*-GFP hetero). Right panel shows the image of the testis of d-rR (recipient) strain. Scale bar, 100 μm. (b) Upper and lower panels show the image of *actb*-GFP hetero and d-rR testis, respectively. Left and middle panels show images of DAPI (blue) and GFP (green), respectively, in the same section; right panel shows the merged image. The GFP signal in germ cells was faint in the fluorescent observation. Scale bar, 50 μm.

## Notes

### Competing Interest Statement

The authors have declared no competing interest.

