## Supplementary figures and images for "Allogeneic testes transplanted into partially castrated adult medaka (*Oryzias latipes*) can produce donor-derived offspring by natural mating over a prolonged period"

### Supplementary Figure 1

#### Supplementary Figure 1

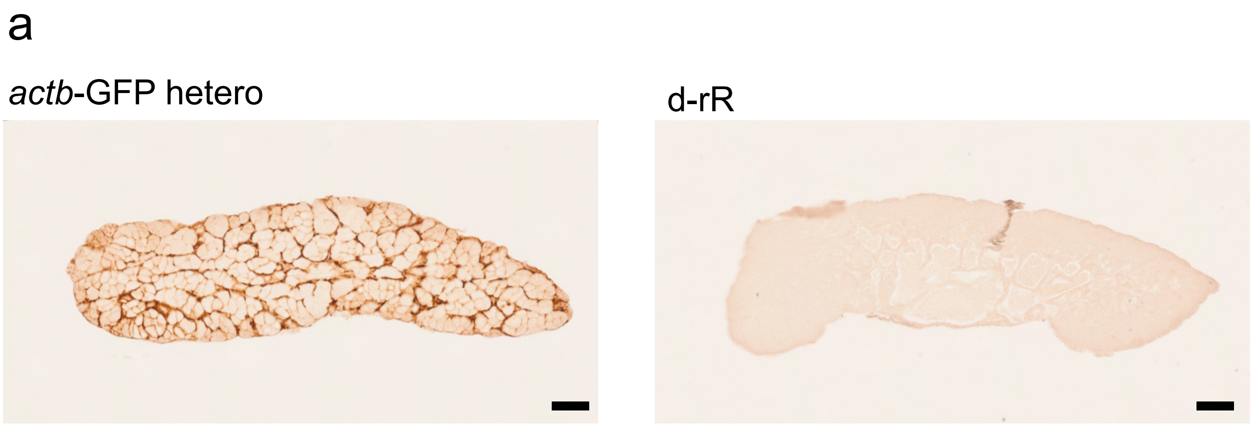


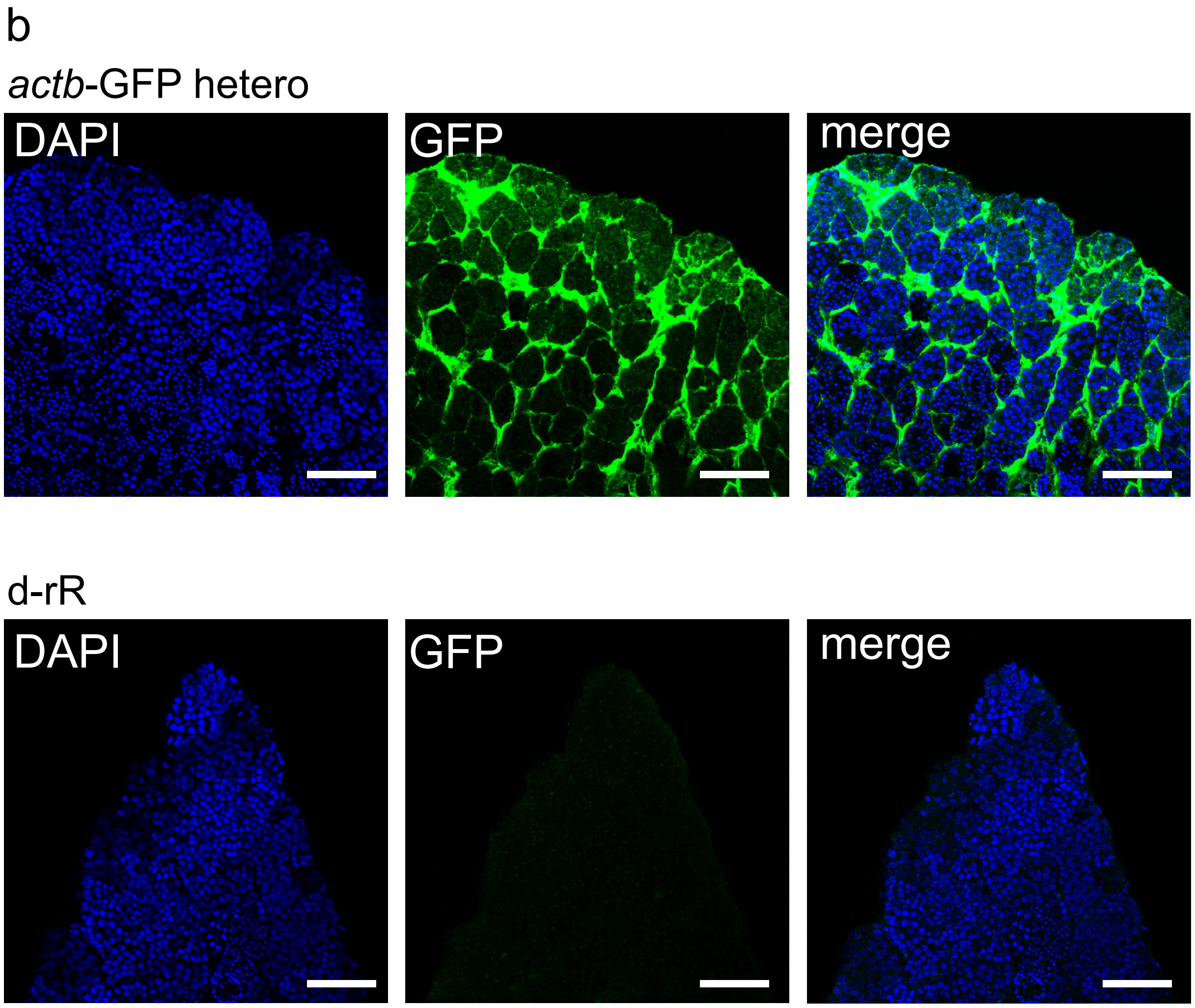
